# Pharmacology of Berberine and its Metabolites, is it the natures Ozempic or Imatinib?

**DOI:** 10.1101/2023.08.05.552100

**Authors:** Naresh Kumar Singh, Muralikrishnan Dhanasekaran, Arun HS Kumar

## Abstract

**Background:** Berberine, which is a naturally occurring alkaloid is widely explored for several health benefits including for weight management and metabolic disorders. The major pharmacological action of berberine is reported to be by activation of AMP-activated protein kinase, while its other clinical outcomes are devoid of clear mechanism of action/s. Hence in this study a detailed pharmacology of berberine and its two major metabolites (berberrubine, and jatrorrhizine) in humans was evaluated using well established Insilco tools.

**Materials and Methods:** The targets of berberine and its metabolites were identified in SwissTargetPrediction server and their affinity was assed using AutoDock vina 1.2.0. The binding pockets of the highest ligand receptor combinations was assessed using the PrankWeb: Ligand Binding Site Prediction tool.

**Results:** Kinases, enzymes and family A GPCR’s were identified as the top three target category of berberine and its metabolites. ROCK2, PIK3CD, KCNMA1, CSF1R and KIT were observed to be the high affinity targets of berberine and its metabolites with affinity values of <4 uM. The affinity of berberine and its metabolites against all AMPKs and lipid/glucose regulator targets (LDLR, DDP4 and PCSK9) were > 10 uM. The IC50 value of berberine and its metabolites against ROCK2 was the least (<1 uM), while their other high affinity targets (PIK3CD, KCNMA1, CSF1R and KIT) showed IC50 values < 5 uM.

**Conclusion:** The diverse range of protein targets and the observed novel high affinity targets (ROCK2, PIK3CD, KCNMA1, CSF1R and KIT) offer valuable insights into the potential mechanisms of action and therapeutic effects of berberine and its metabolites in various disease conditions, which warrants validation in suitable efficacy analysis studies.

## Introduction

Berberine, is a naturally occurring alkaloid compound found in various medicinal plants belonging to Berberidaceae and Ranunculaceae families. Berberis species, such as Berberis aristate, Coptis trifolia, Mahonia bealei and Hydrastis canadensis, has garnered significant attention in recent years due to its diverse pharmacological properties and therapeutic potential.^[1, 2]^ Some common natural source of berberine along with their geographical locations are outlined in table 1. These plants contain varying amounts of berberine in different parts, such as the roots, stems, leaves, or fruits. Berberine is commonly found in the roots and rhizomes of these plants, and it is extracted through various methods for use in traditional medicine and pharmaceutical applications.^[3, 4]^ The berberine content can vary depending on the plant’s species, geographical location, and growing conditions, while the extraction methods and purification processes used play a crucial role in influencing the quality of berberine used for research and medicinal purposes.^[5]^ With a rich history spanning centuries in traditional medicine, berberine has been extensively used in traditional healing practices to address a wide array of ailments. In the pharmaceuticals era, this potent alkaloid has captured the interest of researchers and scientists worldwide, prompting rigorous investigations into its multifaceted pharmacological effects and potential applications in various disease conditions.^[6-10]^ The molecular structure of berberine consists of a quaternary ammonium cation linked to a benzylisoquinoline skeleton, conferring it with unique physicochemical properties and biological activities.^[11]^ The distinctive chemical composition has been associated with a plethora of biological effects, including antioxidant, anti-inflammatory, antimicrobial, antidiabetic, anticancer, and cardiovascular properties, among others.^[1-4]^ These attributes have propelled berberine into the spotlight as a promising natural compound with potential therapeutic benefits for a wide range of medical conditions.

**Table 1:** Common natural source of berberine and their major geographical locations.

| Plants | Geographical Location |
| --- | --- |
| *Berberis vulgaris* (European barberry) | Middle east, Europe |
| *Berberis aristata* (Indian barberry) | Asia |
| *Berberis aquifolium* (Mountain grape) | America |
| *Berberis bealei* (Oregon grape) | America |
| *Berberis japonica* (Beale's barberry) | Asia |
| *Coptis chinensis* (Chinese goldthread) | Asia |
| *Coptis japonica* (Japanese goldthread) | Asia |
| Coptis trifolia (Threeleaf goldthread) | Asia and America |
| Hydrastis canadensis (Goldenseal) | America |
| Mahonia aquifolium (Oregon grape) | America |
| Mahonia bealei (Beale's barberry) | Asia |
| Mahonia fortunei (Fortune's barberry) | Asia |
| Tinospora cordifolia (Heart-leaved moonseed) | Asia |
| Phellodendron amurense (Amur cork tree) | Asia |

The growing body of preclinical and clinical studies on berberine has yielded intriguing results, showing its potential in modulating several key cellular signalling pathways and molecular targets. Berberine exerts its pharmacological effects through multiple mechanisms of action, making it a versatile and promising natural compound with potential therapeutic applications.^[12-14]^ Some of the key mechanisms of action of berberine include: 1) Regulation of Cellular Signalling Pathways: Berberine can modulate various cellular signalling pathways, including AMP-activated protein kinase (AMPK)^[15]^ and mitogen-activated protein kinase (MAPK) pathways.^[16]^ Activation of AMPK helps regulate energy metabolism and glucose homeostasis, making berberine a potential candidate for managing metabolic disorders such as type 2 diabetes. 2) Interaction with Enzymes: Berberine can interact with several enzymes, affecting their activity and function. For instance, it inhibits dipeptidyl peptidase-4 (DPP-4), an enzyme involved in the breakdown of incretin hormones, thus prolonging incretin hormone action and improving insulin secretion in diabetes management.^[17]^ 3) Anti-inflammatory Activity: Berberine has been shown to possess anti-inflammatory properties by inhibiting the production of pro-inflammatory cytokines and reducing the activation of nuclear factor-kappa B (NF-kB).^[18]^ This anti-inflammatory action may contribute to its potential in treating inflammatory conditions. 4) Antioxidant Effects: Berberine has demonstrated antioxidant activity by neutralizing free radicals and reducing oxidative stress. This property can protect cells and tissues from damage caused by reactive oxygen species (ROS) and may have implications for various oxidative stress-related diseases.^[19, 20]^ 5) Antibacterial and Antimicrobial Activity: Berberine exhibits potent antibacterial and antimicrobial properties, making it effective against a wide range of pathogens, including bacteria, viruses, fungi, and parasites. It can inhibit bacterial growth and disrupt the integrity of microbial cell membranes.^[21, 22]^ 6) Modulation of Gut Microbiota: Berberine has the ability to modulate the composition and diversity of the gut microbiota. By promoting the growth of beneficial bacteria and inhibiting harmful species, berberine may contribute to gut health and overall well-being.^[23, 24]^ 7) Anticancer Effects: Berberine has demonstrated anticancer activity by inducing cell cycle arrest, promoting apoptosis (programmed cell death), and inhibiting the growth and metastasis of cancer cells. It also exhibits anti-angiogenic properties, hindering the development of new blood vessels that support tumour growth.^[25, 26]^ 8) Cholesterol and Lipid Regulation: Berberine can lower cholesterol levels by inhibiting the enzyme HMG-CoA reductase, which is involved in cholesterol synthesis. It also enhances the expression of LDL receptors, promoting the clearance of LDL cholesterol from the bloodstream.^[27]^ 9) Neuroprotective Effects: Berberine has been investigated for its potential neuroprotective properties. It can modulate neurotransmitter systems and attenuate neuroinflammation, offering potential benefits in neurodegenerative diseases and cognitive disorders.^[28]^ Despite these diverse mechanisms of actions, It is important to note that the exact mechanisms of action of berberine may vary depending on the specific cellular context and the target tissue or organ. Additionally, more research is needed to fully elucidate the comprehensive mechanisms underlying berberine’s diverse pharmacological effects. Nevertheless, the multifaceted nature of berberine’s mechanisms of action contributes to its wide range of therapeutic potentials, making it an intriguing candidate for further investigation in various disease conditions. Despite the increasing evidence supporting the pharmacological activities of berberine, several critical questions remain unanswered. The mechanisms governing its multitargeted effects, together with its pharmacokinetic properties, and the factors influencing its bioavailability are areas that warrant further exploration. Furthermore, while numerous clinical trials have demonstrated the therapeutic potential of berberine in certain disease conditions, additional robust clinical investigations are necessary to elucidate its safety and efficacy profile across different patient populations.

In recent years, advancements in systems biology and computational approaches have revolutionized drug discovery, enabling researchers to comprehend complex interactions within biological systems. Network analysis, a powerful tool in this domain, has gained prominence as an effective method to unravel the intricacies of pharmacological actions and reveal the underlying mechanisms of active compounds like berberine. Network analysis offers a comprehensive and holistic perspective by representing biological entities, such as proteins, genes, or metabolites, as nodes and their interactions as edges in a network.^[29-31]^ This approach not only elucidates individual components’ roles but also emphasizes the importance of interconnectedness and crosstalk within cellular pathways. The integration of diverse omics data, including genomics, proteomics, and metabolomics, in conjunction with network analysis enables the construction of sophisticated networks that can capture the multifaceted interactions involved in berberine’s pharmacological effects. While previous studies have shed light on some of the molecular targets and signalling pathways impacted by berberine, a comprehensive systems-level analysis of its pharmacological mechanisms remains limited. The utilization of network analysis in investigating the pharmacology of berberine provides a unique opportunity to unravel the complexity of its mode of action, offering valuable insights into its therapeutic potential and possible adverse effects. In this study, a comprehensive examination of the pharmacology of berberine was performed using the Insilco tools, aiming to shed light on the molecular mechanisms underlying its multifaceted effects. The findings from this study will not only enhance our understanding of berberine’s biological activities but also provide valuable insights that could contribute to the development of novel therapeutic strategies harnessing the potential of this interesting herbal alkaloid.

## Materials and Methods

The isomeric SMILES sequence of berberine and its metabolites (berberrubine, and jatrorrhizine) obtained from the PubChem database were inputted into the SwissTargetPrediction server to identify the targets specific to homo sapiens. The affinity values of berberine and its metabolites with their respective targets was assessed using AutoDock vina 1.2.0 as reported before for other ligand-receptor combinations.^[30-32]^

The top 10 target network of berberine was identified from the STITCH database (https://stitch-db.org) and the affinity of berberine and its metabolites with these targets was assessed using AutoDock vina 1.2.0. As AMPK is reported to be a major target of berberine, the network protein analysis of human AMPK was conducted as reported before using the STRING Database (https://string-db.org), and the affinity of berberine and its metabolites against all known forms of human AMPK identified in this network was evaluated using AutoDock vina 1.2.0. In addition some selective lipid targets (GLP1R, ZGLP1, DPP4; Uniprot ID P43220, P0C6A0, P27487 respectively) and targets of berberine listed in the DrugBank (BIRC5 and qacR; Uniprot ID O15392, P0A0N5 respectively) were also assessed for affinity using AutoDock vina 1.2.0.^[30-32]^

The pharmacokinetic parameters of berberine and its metabolites was assessed using the SwissADME server. The targets with top five affinity values with berberine or its metabolites were further assessed using the PrankWeb: Ligand Binding Site Prediction tool (https://prankweb.cz/) to identify the major binding sites and an inhouse algorithm reported previously was used to estimate the IC_50_ values of berberine and its metabolites against each of the top five targets.

## Results

Berberine belongs to the class of benzylisoquinoline alkaloids and its chemical structure can be divided into three main parts: the quaternary ammonium group, the benzylisoquinoline core, and the methoxy group (Figure 1). The quaternary ammonium group (-NR4+) in berberine contains four carbon atoms bonded to a central nitrogen atom. It confers a positive charge to the nitrogen atom, making berberine a water-soluble compound. The benzylisoquinoline core of berberine consists of two aromatic rings (A and B) connected by a bridgehead carbon (C). Ring A is a quaternary benzene ring, and ring B is a partially hydrogenated aromatic ring. The bridgehead carbon (C) is a unique feature of the benzylisoquinoline alkaloids, and in berberine, it plays a crucial role in the subsequent formation of its metabolites. At position C9 of the B-ring, berberine contains a methoxy group (OCH3). This functional group is important for understanding how berberine is converted to its metabolites. Berberine is metabolized in the body through various enzymatic reactions, leading to the formation of several metabolites. Two significant metabolites of berberine are berberrubine and jatrorrhizine (Figure 1). The conversion of berberine to berberrubine involves the oxidation of the methoxy group (OCH3) at position C9 of the B-ring. This oxidation reaction replaces the methoxy group with a hydroxy group (OH), resulting in the formation of berberrubine. This process is typically facilitated by enzymes in the liver, where most of the metabolism of berberine occurs. While jatrorrhizine is formed through a demethylation reaction, where the methoxy group (OCH3) at position C9 of the B-ring is removed. The demethylation process exposes a hydrogen atom, converting berberine into jatrorrhizine. Like the conversion to berberrubine, this reaction is also catalysed by specific enzymes in the body.

**Figure 1:**
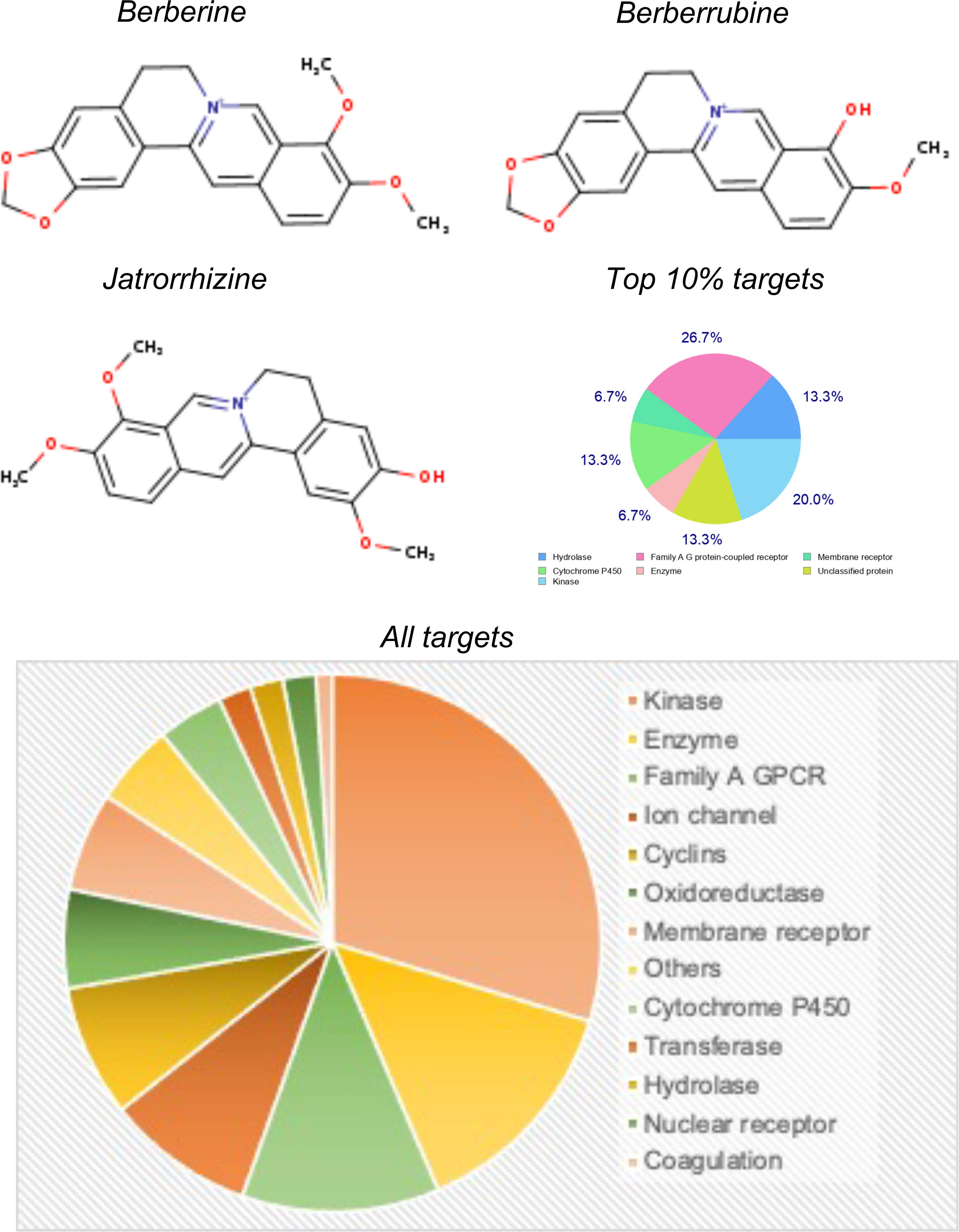
Chemical structure of berberine and its two major metabolites along with their major target categories.

The distribution of different protein targets of berberine and its metabolites is shown in figure 1. There were 29 kinases targeted by berberine and its metabolites. Kinases are enzymes that transfer phosphate groups to other proteins, regulating various cellular processes, including signal transduction, cell cycle control, and metabolism. 14 enzymes were targeted by berberine and its metabolites which are biological catalysts that facilitate biochemical reactions in cells. 12 Family A G Protein-Coupled Receptors (GPCRs) were targeted by berberine and its metabolites. GPCRs are a large family of membrane receptors that play crucial roles in cell signalling and are involved in various physiological processes. 9 ion channels were targeted by berberine and its metabolites, which are integral membrane proteins that allow the passage of ions across cell membranes, controlling electrical signalling in cells. 8 cyclins were targeted by berberine and its metabolites, which are proteins that regulate the cell cycle. 7 types of membrane receptors were targeted by berberine and its metabolites, which are proteins located on the cell membrane that interact with extracellular molecules to initiate cellular responses. 6 oxidoreductases were targeted by berberine and its metabolites, which are a class of enzymes involved in redox reactions. 4 cytochrome P450 enzymes were targeted by berberine and its metabolites, which are a superfamily of enzymes involved in the metabolism of various compounds, including drugs and toxins. 2 transferases were targeted by berberine and its metabolites, which are a class of enzymes that transfer functional groups between molecules. 2 compounds hydrolases were targeted by berberine and its metabolites, which are enzymes that catalyse hydrolysis reactions, breaking down compounds with the addition of water. 2 nuclear receptors were targeted by berberine and its metabolites, which are a class of ligand-activated transcription factors that regulate gene expression. 1 protein involved in the coagulation cascade was targeted by berberine and its metabolites, which is critical to the process of blood clot formation. A diverse range of protein targets were targeted by berberine and its metabolites, with a significant preference towards kinases, enzymes and family A GPCR’s. Kinases, enzymes and family A GPCR’s were the top three category of targets of berberine and its metabolites (Figure 1). Affinity analysis of berberine and its metabolites against each of its identified targets in humans suggested that ROCK2, PIK3CD, KCNMA1, CSF1R and KIT were the high affinity targets with affinity values of <4 uM (Figure 2, tables 2, 3 and 4). A complete list of all targets of berberine and its metabolites is shown in figure 2, while the targets with affinity values <10 uM are shown in tables 2, 3 and 4. The affinity of berberine and its metabolites with their targets ranged from 20.98 uM to 450 uM (Figure 2).

**Figure 2:**
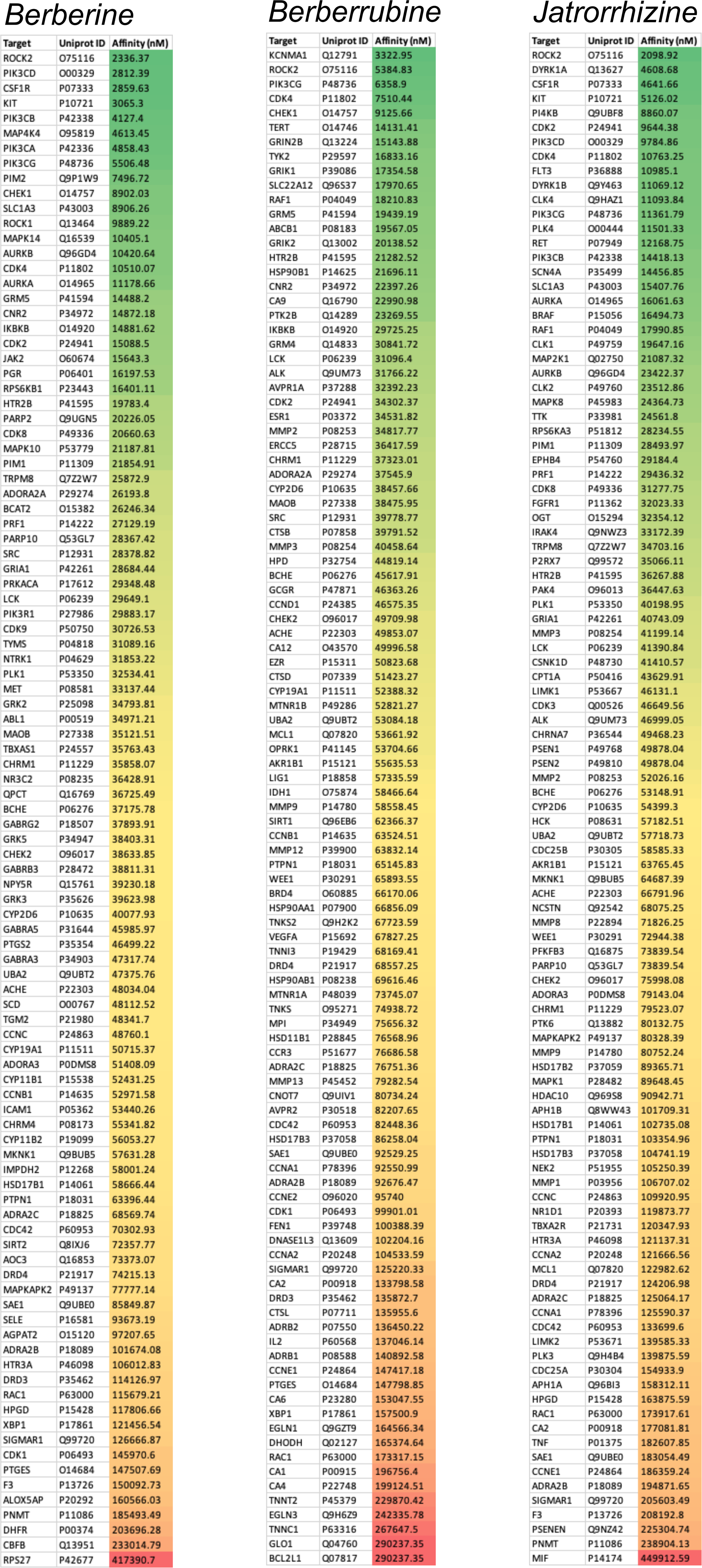
Affinity values of berberine and its two major metabolites with all their targets in humans identified in SwissTargetPrediction server. The affinity values are arranged in ascending order and colour coded (green with highest affinity and red with lowest affinity values).

**Table 2:** High affinity targets of berberine.

| Target | Gene | Uniprot ID | Affinity (nM) | Target Class |
| --- | --- | --- | --- | --- |
| Rho-associated protein kinase 2 | ROCK2 | O75116 | 2336.4 | Kinase |
| PI3-kinase p110-delta subunit | PIK3CD | O00329 | 2812.4 | Enzyme |
| Macrophage CSF receptor | CSF1R | P07333 | 2859.6 | Receptor |
| Stem cell growth factor receptor | KIT | P10721 | 3065.3 | Receptor |
| PI3-kinase p110-beta subunit | PIK3CB | P42338 | 4127.4 | Enzyme |
| Mitogen-activated protein kinase 4 | MAP4K4 | O95819 | 4613.5 | Kinase |
| PI3-kinase p110-catalytic | PIK3CA | P42336 | 4858.4 | Enzyme |
| PI3-kinase p110-gamma subunit | PIK3CG | P48736 | 5506.5 | Enzyme |
| STPK PIM2 | PIM2 | Q9P1W9 | 7496.7 | Kinase |
| STPK Chk1 | CHEK1 | O14757 | 8902.0 | Kinase |
| Excitatory amino acid transporter 1 | SLC1A3 | P43003 | 8906.3 | Ion channel |
| Rho-associated protein kinase 1 | ROCK1 | Q13464 | 9889.2 | Kinase |
| MAP kinase p38 alpha | MAPK14 | Q16539 | 10405.1 | Kinase |
| STPK Aurora-B | AURKB | Q96GD4 | 10420.6 | Kinase |
| Cyclin-dependent kinase 4 | CDK4 | P11802 | 10510.1 | Kinase |

**Table 3:** High affinity targets of berberrubine.

| Target | Gene | Uniprot ID | Affinity (nM) | Target Class |
| --- | --- | --- | --- | --- |
| Calcium-activated potassium channel subunit alpha-1 | KCNMA1 | Q12791 | 3322.95 | Ion channel |
| Rho-associated protein kinase 2 | ROCK2 | O75116 | 5384.83 | Kinase |
| PI3-kinase p110-gamma subunit | PIK3CG | P48736 | 6358.9 | Enzyme |
| Cyclin-dependent kinase 4 | CDK4 | P11802 | 7510.44 | Cyclins |
| STPK Chk1 | CHEK1 | O14757 | 9125.66 | Kinase |

**Table 4:** High affinity targets of jatrorrhizine.

| Target | Gene | Uniprot ID | Affinity (nM) | Target Class |
| --- | --- | --- | --- | --- |
| Rho-associated protein kinase 2 | ROCK2 | O75116 | 2098.92 | Kinase |
| Dual-specificity tyrosine-phosphorylation regulated kinase 1A | DYRK1A | Q13627 | 4608.68 | Kinase |
| MCSF receptor | CSF1R | P07333 | 4641.66 | Receptor |
| Stem cell growth factor receptor | KIT | P10721 | 5126.02 | Receptor |
| PI4-kinase beta subunit | PI4KB | Q9UBF8 | 8860.07 | Kinase |
| Cyclin-dependent kinase 2 | CDK2 | P24941 | 9644.38 | Cyclins |
| PI3-kinase p110-delta subunit | PIK3CD | O00329 | 9784.86 | Kinase |
| Cyclin-dependent kinase 4 | CDK4 | P11802 | 10763.25 | Kinase |
| Tyrosine-protein kinase receptor | FLT3 | P36888 | 10985.1 | Kinase |

The top ten targets of berberine and its metabolites identified in the STITCH database were also subjected to the affinity analysis. Among these the top three targets were the ones associated with lipid regulation i.e., LDLR, DDP4 and PCSK9 (Figure 3). However all these targets showed affinity values > 10 uM. Additionally the affinity of berberine against GLP1 (Uniprot ID P0C6A0) and its receptor (Uniprot ID P43220) was observed to be 75.29 uM and 45.44 uM respectively. BIRC5 (Uniprot ID O15392) and qacR (Uniprot ID P0A0N5) are also reported to be targets of berberine in the drug bank database and their affinity was observed to be 39.08 uM and 23.94 uM respectively. Inhibition of AMPK has been reported to be the major mechanism of action by which berberine is reported to exhibit its pharmacodynamic effects. Hence in this study all known forms of human AMPK were assessed for its affinity with berberine, which ranged from 12 to 104 uM (figure 3). Among the human AMPKs, PRKAA2 showed the highest affinity of 12.03 uM while the PRKAB2 had the least affinity of 104.12 uM (Figure 3).

**Figure 3:**
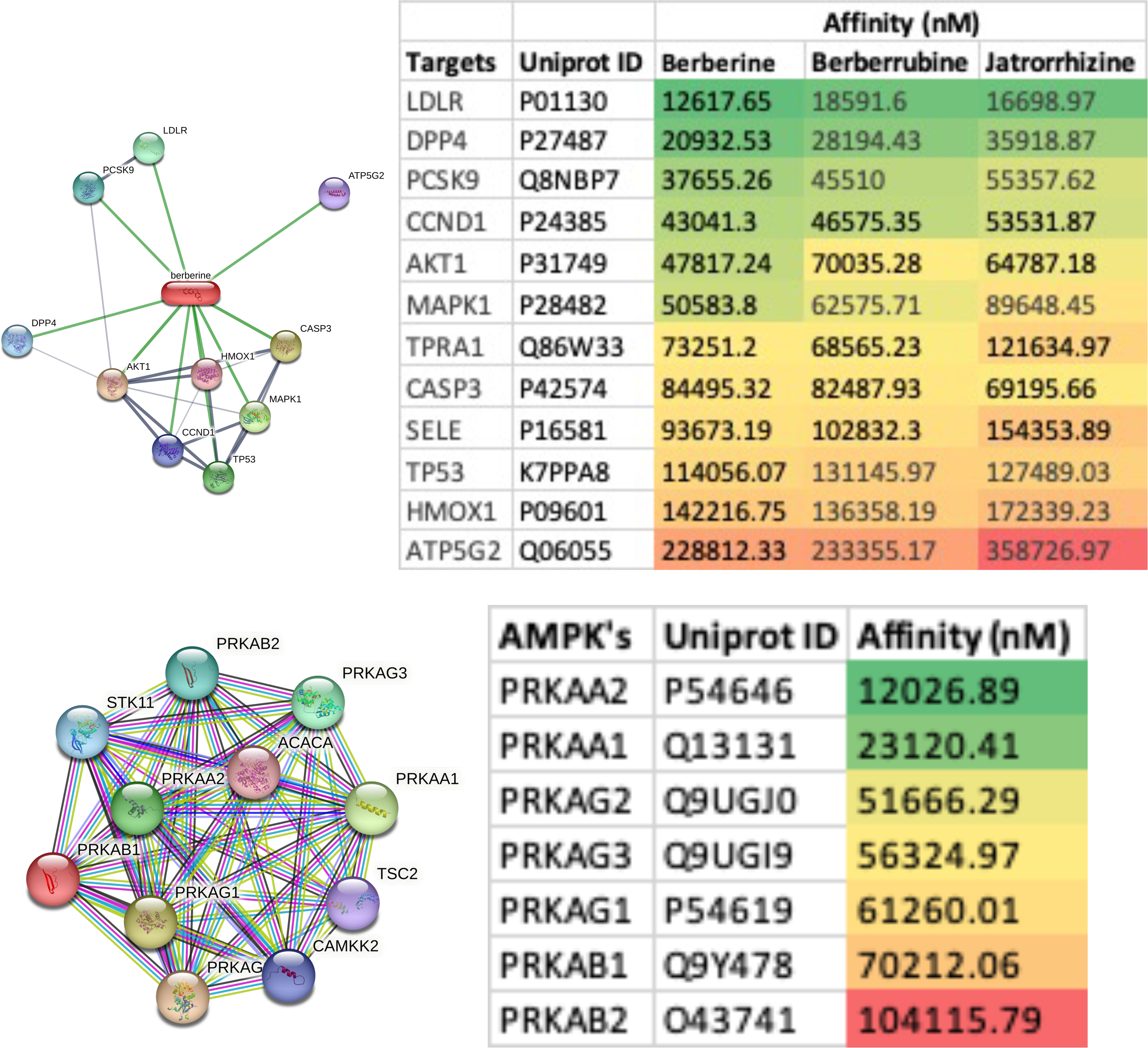
Affinity values of berberine and its two major metabolites with their target networks (Network image) identified in the STITCH database. Bottom panel shows the networks of human AMP-activated protein kinases (AMPK) identified in STRING database along with the affinity values of berberine against all forms of AMPK in humans. The affinity values are arranged in ascending order and colour coded (green with highest affinity and red with lowest affinity values).

A detailed ADME analysis of berberine and its metabolites was performed using the SwissADME database and the relevant Physicochemical Properties, Lipophilicity, solubility, Pharmacokinetics, Druglikeness parameters are summarised in table 5 and figure 4. The binding affinity of berberine and its metabolites with their high affinity targets was performed using AutoDock vina to identify the binding pockets and their IC_50_ values, which are summarised in figure 4. The least IC_50_ value of berberine and its metabolites was for ROCK2 (<1 uM). The details of the best binding pocket for each of the high affinity ligand-receptor combinations is summarised in table 6.

**Figure 4:**
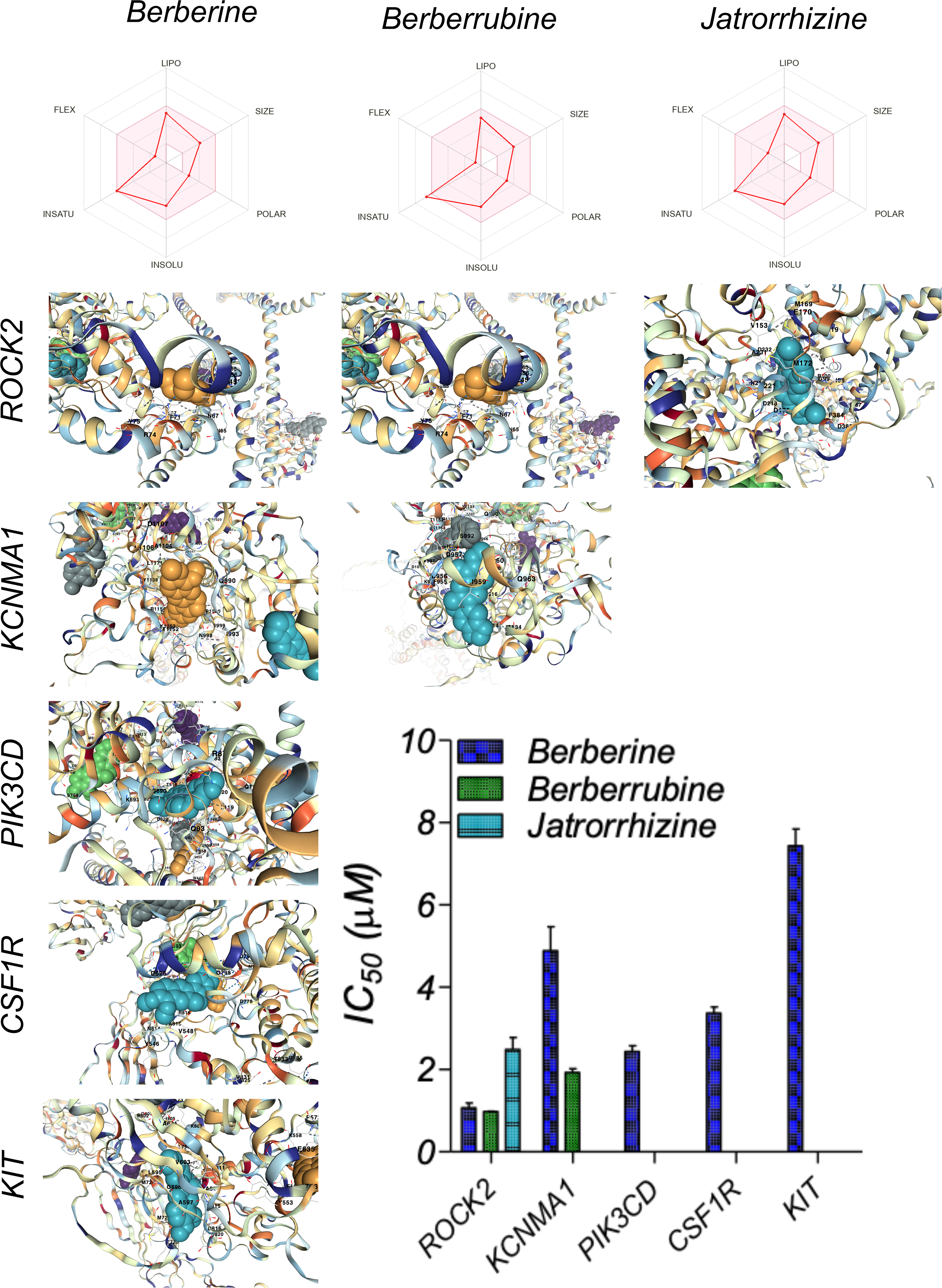
Pharmacokinetic parameters of berberine and its two major metabolites. The binding pockets of high affinity ligand-receptor combinations of berberine and its two major metabolites with their respective targets is shown along with its IC_50_ values (bar graph).

**Table 5:** Physicochemical Properties, Lipophilicity, Solubility, Pharmacokinetics, and Druglikeness parameters of berberine and its two major metabolites.

|  | Berberine | Berberrubine | Jatrorrhizine |
| --- | --- | --- | --- |
| Mol Wt | 336.36 | 322.33 | 338.38 |
| Heavy atoms | 25 | 24 | 25 |
| Aromatic heavy atoms | 16 | 16 | 16 |
| Fraction Csp3 | 0.25 | 0.21 | 0.25 |
| Rotatable bonds | 2 | 1 | 3 |
| H-bond acceptors | 4 | 4 | 4 |
| H-bond donors | 0 | 1 | 1 |
| Molar Refractivity | 94.87 | 90.41 | 97.33 |
| TPSA (Surface area) | 40.8 | 51.8 | 51.8 |
| Log<sub>P</sub> | 3.74 | 3.21 | 3.43 |
| Consensus Log P | 2.53 | 2.24 | 2.31 |
| Log<sub>S</sub> | -4.16 | -4.05 | -4.19 |
| Solubility (mg/ml) | 2.30E-02 | 2.85E-02 | 2.20E-02 |
| Solubility (mol/l) | 6.85E-05 | 8.85E-05 | 6.49E-05 |
| Class | Moderately soluble | Moderately soluble | Moderately soluble |
| Log<sub>Sw</sub> | -5.92 | -5.23 | -5.72 |
| GI absorption | High (~97%) | High (~97%) | High (~97%) |
| BBB permeant | Yes | Yes | Yes |
| Pgp substrate | Yes | Yes | Yes |
| CYP1A2 inhibitor | Yes | Yes | Yes |
| CYP2C19 inhibitor | No | No | No |
| CYP2C9 inhibitor | No | No | No |
| CYP2D6 inhibitor | Yes | Yes | Yes |
| CYP3A4 inhibitor | Yes | Yes | Yes |
| log<sub>Kp</sub> (cm/s) | -5.78 | -5.93 | -5.94 |
| Lipinski violations | 0 | 0 | 0 |
| Ghose violations | 0 | 0 | 0 |
| Veber violations | 0 | 0 | 0 |
| Egan violations | 0 | 0 | 0 |
| Muegge violations | 0 | 0 | 0 |
| Bioavailability Score | 0.55 | 0.55 | 0.55 |
| PAINS #alerts | 0 | 0 | 0 |
| Brenk alerts | 1 | 1 | 1 |
| Lead likeness violations | 1 | 0 | 0 |
| Synthetic Accessibility | 3.14 | 3.01 | 3.06 |
| Plasma Protein Binding | 58.54 | 58.54 | 58.54 |

**Table 6:**
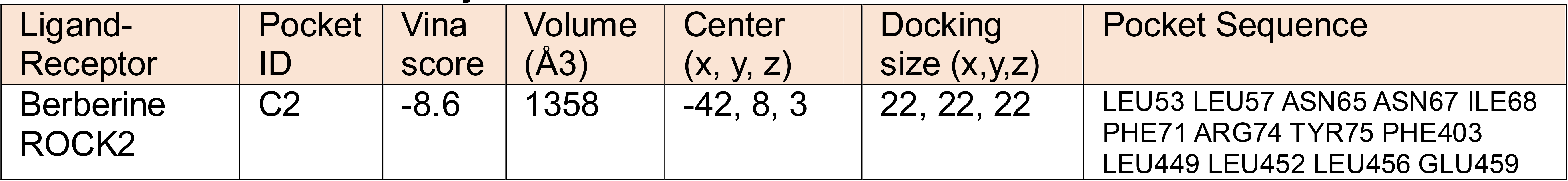
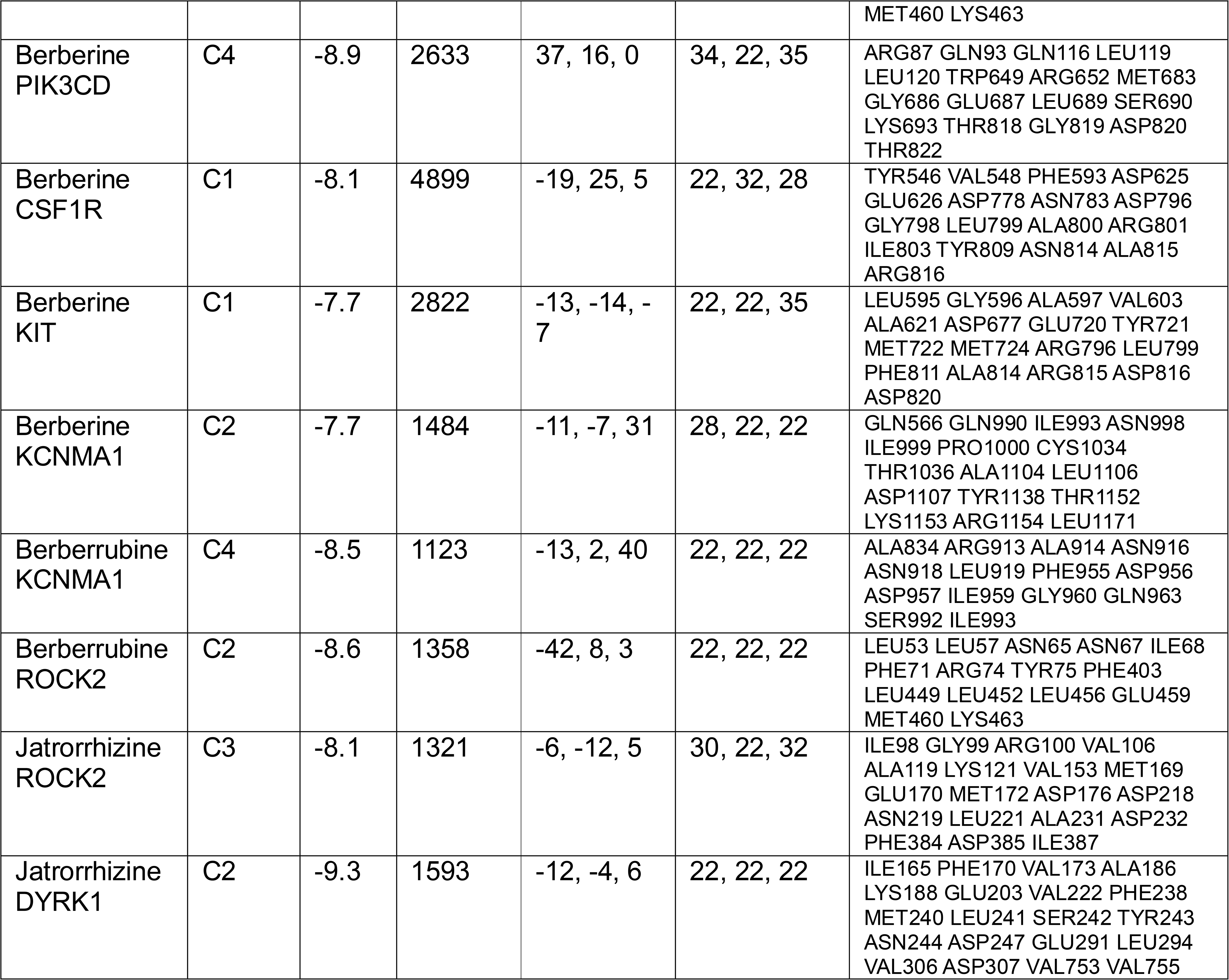
The binding pockets of high affinity ligand-receptor combinations of berberine and its two major metabolites.

## Discussion

The results from this study indicate that berberine and its metabolites have a diverse range of protein targets, and the top three categories of targets are kinases, enzymes, and Family A G Protein-Coupled Receptors (GPCRs). This aligns with existing literature on the pharmacology of berberine and its metabolites.^[2, 24]^ The significant targeting of kinases by berberine and its metabolites is in line with previous research that highlights the importance of kinase inhibition as a major mechanism of action for berberine. Kinases play crucial roles in signal transduction pathways, and their inhibition can lead to various physiological effects, including anti-inflammatory,^[33]^ anticancer,^[34]^ and antidiabetic^[35]^ properties. Among the kinases AMPK (AMP-activated protein kinase) is extensively reported to be major target of berberine pharmacodynamics as AMPK activation is considered a major mechanism by which berberine exerts its beneficial effects.^[36, 37]^ AMPK activation leads to increased glucose uptake, improved insulin sensitivity, and enhanced fatty acid oxidation, contributing to its potential antidiabetic and metabolic effects. However the affinity analysis of berberine against various forms of human AMPK showed values ranging from 12 to 104 uM, which was much higher than the affinity of berberine and its metabolites against its high affinity targets (ROCK2, PIK3CD, KCNMA1, CSF1R and KIT), for which the affinity values were <4 uM. PRKAA2 showed the highest affinity of 12.03 uM, while the affinity with rest of the AMPK’s was >20 uM. Hence it is reasonable to assume that influence of berberine on AMPK physiology is regulated by activation of PRKAA2, although the dose of berberine required to achieve this will be high. PRKAA2, which is highly expressed in kidneys, heart and skeletal muscles, plays a crucial role in coordinating cellular responses to fluctuations in energy levels, making it a central player in maintaining cellular energy homeostasis and metabolic regulation. Its functions have implications for various metabolic diseases and potential therapeutic interventions for metabolic disorders, such as type 2 diabetes, obesity, and cardiovascular diseases.^[38, 39]^

Berberine is also reported to target various enzymes involved in cellular processes. For example, it has been reported to inhibit the enzyme dipeptidyl peptidase-4 (DPP4),^[17]^ which plays a role in glucose regulation, and this could contribute to its antidiabetic effects. The regulation of other related targets may also contribute to the diverse therapeutic potential of berberine in metabolic and lipid disorders. Hence it wasn’t surprising to find the top targets (LDLR, DPP4, and PCSK9) of berberine in the STITCH database were all associated with lipid regulation, which align with berberine’s known effects on lipid metabolism.^[27, 40]^ However all these targets showed affinity values > 10 uM, which is achievable only at higher doses of berberine. It may be likely that berberine underdoes significant bioconcentration (accumulation in tissue niche), which may account for its therapeutics benefits through influencing these high affinity targets. However this remains to be validated in suitable in vivo studies. Despite low bioavailability, berberine is reported to have high tissue distribution. Following oral administration, the levels of berberine and its active metabolites in various organs are reported to be higher than in the bloodstream.^[2]^ The organ distribution of berberine is specifically reported for liver (hepatic metabolism), kidneys (renal excretion), skeletal muscle, lungs, brain, heart, and pancreas. In addition berberine is also reported to remain stable in adipose tissue for over 48-hours, which suggests depot effect in fat tissue, influencing the pharmacokinetics and pharmacodynamics of berberine.^[2]^ The observed organ distribution of berberine highlights its remarkable ability to reach various target sites throughout the body, potentially influencing multiple physiological processes.^[2]^ Understanding the specific distribution patterns of berberine in different organs provides valuable insights into its pharmacological effects and lays the groundwork for optimizing its therapeutic applications in the treatment of various diseases. Further studies investigating the mechanisms and factors influencing the organ distribution of berberine are crucial for harnessing its full therapeutic potential and enhancing our understanding of its pharmacokinetic profile.

In contrast to the known and widely reported targets of berberine, in this study ROCK2, PIK3CD, KCNMA1, CSF1R and KIT were identified as high affinity (<4 uM) targets of berberine and its metabolites. The targetability of these targets were further validated in this study by revealing the specific binding pockets of berberine or its metabolites on these targets at therapeutically feasible concentrations. ROCK2’s involvement in actin cytoskeleton regulation and cell contractility makes it an essential player in various cellular activities, including cell migration, adhesion, proliferation, and tissue development. Dysregulation of ROCK2 has been associated with several diseases, making it a potential target for therapeutic interventions in conditions such as cancer, cardiovascular diseases, and neurological disorders.^[41-44]^ The major pathological role of ROCK2 is its involvement in promoting various aspects of cancer progression and metastasis.^[42, 43]^ Aberrant activation or overexpression of ROCK2 has been associated with cancer development and is linked to several pathological processes in cancer cells. Hence berberine by inhibiting ROCK2 has a vital role as an anticancer therapeutic. The major pathological role of PIK3CD is its association with immunodeficiency disorders caused by mutations in the PIK3CD gene.^[45, 46]^ These disorders are collectively known as “PIK3CD-related primary immunodeficiency” or “Activated PI3K Delta Syndrome” (APDS).^[47, 48]^ APDS is characterized by dysregulation of the PI3K signalling pathway, leading to aberrant immune cell function and impaired immune responses. There are two types of APDS: APDS1 and APDS2, depending on the specific genetic mutation involved. The potential of berberine and its metabolites to inhibit PIK3CD with high affinity offers a viable options in the treatment of APDS for which currently very limited precision medicine options are available.

KCNMA1 which was also an high affinity target of berberine and its metabolites is also known as BKCa (Big Potassium Calcium-Activated Channel), is associated with various cardiovascular and neurological disorders.^[49, 50]^ KCNMA1 encodes the alpha-subunit of the BKCa channel, which is a large-conductance, calcium-activated potassium channel. Dysregulation or mutations in the KCNMA1 gene can lead to the cardiovascular disorders such as hypertension, arrythmias and vasospasms and following neurological disorders, epilepsy, Parkinson’s disease and dementia.^[51]^ Also overexpression of BKCa channels has been associated with increased invasion and metastasis in certain types of cancer.^[52-54]^ In addition abnormal BKCa channel function is also reported to be associated with gastrointestinal motility disorders and urinary incontinence. It remains to be established how berberine and its metabolites can be therapeutically beneficial for these clinical conditions directly associated with abnormal BKCa channel function.

CSF1R is a receptor tyrosine kinase that binds to Colony-Stimulating Factor 1 (CSF-1 or M-CSF), a cytokine involved in regulating the differentiation, survival, and function of macrophages and other myeloid cells and is various diseases and is primarily associated with dysregulated immune responses and abnormal cell proliferation. Dysregulation of CSF1R signalling has been implicated in various cancers, particularly those of myeloid cell origin.^[55, 56]^ Overexpression or activation of CSF1R in cancer cells or in the tumour microenvironment can lead to increased recruitment, proliferation, and survival of tumour-associated macrophages (TAMs).^[57]^TAMs play a critical role in promoting tumour progression by creating an immunosuppressive microenvironment, promoting angiogenesis, and facilitating tumour invasion and metastasis. Targeting CSF1R signalling has emerged as a potential therapeutic strategy for cancer treatment and paraphs berberine will prove to be a valuable therapeutic as anticancer agent due to its high affinity against CSF1R. CSF1R signalling is involved in the regulation of macrophage and monocyte functions, which play a central role in the immune response to infection and inflammation. Dysregulated CSF1R signalling can lead to excessive or aberrant activation of macrophages, contributing to chronic inflammatory conditions such as rheumatoid arthritis, atherosclerosis, unstable plaques and inflammatory bowel diseases.^[58, 59]^ However CSF1R is also involved in several physiological process including differentiation and function of osteoclasts, microglia’s, macrophages and monocytes. Given the diverse roles of CSF1R in regulating immune responses, cell proliferation, and tissue homeostasis, its dysregulation can have far-reaching effects on various physiological processes. Understanding the pathological roles of CSF1R is crucial for the development of targeted therapies aimed at modulating its signalling pathways for therapeutic benefit in specific diseases.

KIT is a receptor tyrosine kinase that binds to stem cell factor (SCF), also known as KIT ligand. It plays essential roles in regulating cell survival, proliferation, and differentiation of various cell types. The major pathological role of KIT is its association with several diseases, particularly cancer and certain haematological and gastrointestinal disorders.^[60, 61]^ Gastrointestinal Stromal Tumors (GIST) are the most common mesenchymal tumours of the gastrointestinal tract. The majority of GISTs have activating mutations in the KIT gene, leading to constitutive activation of the KIT receptor.^[62]^ This abnormal activation drives uncontrolled cell proliferation and tumour growth. Although KIT inhibitors, such as imatinib,^[63]^ has shown significant clinical benefit for patients with GISTs, there remains a merit in co-use of imatinib with berberine to achieve therapeutic synergy. Systemic mastocytosis is a rare disorder characterized by an abnormal accumulation of mast cells in various tissues and organs. The majority of cases of systemic mastocytosis have activating mutations in the KIT gene, leading to increased proliferation and survival of mast cells.^[64]^ This results in the release of various mediators, causing symptoms such as skin rashes, itching, flushing, and potentially life-threatening allergic reactions. A similar response is also observed in cytokine release syndrome^[65]^ and the role of KIT in this remains unknown. Nevertheless the higher affinity of berberine against KIT offers a value therapeutic option, which warrants to be investigated. Acute Myeloid Leukemia (AML), some melanomas and seminomas are reported to involve mutations in the KIT gene.^[66, 67]^ These mutations are associated with poor prognosis and resistance to standard chemotherapy. Targeting these KIT mutations with specific inhibitors is an area of ongoing research and berberine could be potentially useful.

The ADME analysis provides insight into the drug-like properties of berberine and its metabolites. Understanding these properties is crucial for assessing their potential for drug development. The reported physicochemical properties, solubility, and pharmacokinetic parameters are important factors in determining the drug-likeness of a compound, which were consistent with several studies reporting drug-likeness characteristics of berberine and its metabolites.^[68-70]^ The drug-likeness of berberine and its metabolites is further validated by the binding affinity analysis and IC_50_ values which provide assurance about the strength of interaction between berberine and its metabolites with their high-affinity targets at therapeutically feasible concentrations.

In summary, this study while aligning well with existing literature on the pharmacology of berberine and its metabolites provides a novel insight into its potential mechanism of actions. The diverse range of protein targets and the observed high affinity targets (ROCK2, PIK3CD, KCNMA1, CSF1R and KIT) offer valuable insights into the potential mechanisms of action and therapeutic effects of berberine and its metabolites in various disease conditions, which warrants validation in suitable efficacy analysis studies. However considering kinases being the major target category and cancer being the major therapeutic category of berberine, it appears to me that berberine is more of a natures Imatinib rather than Ozempic.

## Acknowledgements

Research support from University College Dublin-Seed funding/Output Based Research Support Scheme (R19862, 2019), Royal Society-UK (IES\R2\181067, 2018) and Stemcology (STGY2917, 2022) is acknowledged.

